# Application of fluorescence lifetimes to multi-imaging analysis in plant cells

**DOI:** 10.1101/2023.08.28.555227

**Authors:** Tsuyoshi Aoyama, Nagisa Sugimoto, Yoshikatsu Sato

## Abstract

Multi-imaging analysis has become an indispensable technique to visualize multiple target proteins and intracellular components simultaneously. While current multi-imaging analysis relies on the differences in emission spectra of fluorescent molecules, the use of fluorescence lifetime imaging microscopy (FLIM), which exploits the differences in fluorescence lifetimes of fluorescent proteins, in multi-imaging analysis is quite limited. In this study, we successfully discriminated fluorescent proteins with similar colors but different fluorescence lifetimes *in vitro* and *in planta*. We found that four fluorescent proteins with similar emission spectra could be distinguished by FLIM. In addition, we found that FLIM could clearly separate fluorescent proteins if they differ by at least 0.2 ns. In a proof-of-concept experiments for plant live imaging, we transiently expressed fluorescent proteins with different subcellular localization tags in *Physcomitrium patens* by particle bombardment. Each fluorescent protein exhibited its fluorescence lifetime at the subcellular localization corresponding to the localization tag in *P. patens* with little or no effect of chlorophyll autofluorescence. Our results demonstrate the effectiveness of FLIM in revealing the spatiotemporal dynamics of a large number of fluorescent proteins in living plant cells.

## Introduction

Fluorescent proteins, which are available in a variety of color options, have become an indispensable tool for multi-imaging analysis to analyze complex biological phenomena (Müller-Taubenberger and Anderson, 2007). Combining fluorescent proteins with different emission spectra is widely used to visualize multiple target proteins and intracellular components simultaneously (Heim et al., 1994; Matz et al., 1999; Lin et al., 2009; Filonov et al., 2011; Ormo et al., 1996). However, relying solely on the emission spectra (color) and its brightness-based separation limits the number of fluorescent molecules that can be simultaneously identified.

Fluorescence lifetimes are defined as the time a molecule spends in the excited state before returning to the ground state and are generally on the order of nanoseconds (ns). This parameter is an intrinsic value of a fluorescent molecule and is an important factor in the properties of fluorophores, including fluorescent proteins. Fluorescent proteins with various fluorescence lifetimes as well as emission spectra have been developed, with fluorescence lifetimes ranging from 0.16 to 5.1 ns (Murakoshi et al., 2015; Sarkisyan et al., 2015; Mamontova et al., 2018; Mukherjee et al., 2022).

Fluorescence intensity-based microscopy can distinguish between fluorescent molecules with different emission spectra but cannot distinguish between fluorescent molecules with similar spectra. However, fluorescence lifetime imaging microscopy (FLIM) can be used to identify fluorescent proteins based on their fluorescence lifetime rather than their emission spectra (Valeur, 2001). Furthermore, among the photophysical properties of molecules, the fluorescence lifetime is highly dependent on the environment around the fluorescent molecule. This property has led to the development of fluorescent molecular sensors to monitor various intracellular environments, such as temperature, pH, ion concentration, and DNA density (Hille et al., 2009; Ogikubo et al., 2011; Okabe et al., 2012; Uno et al., 2021).

FLIM is also available for Förster resonance energy transfer (FRET) experiments to detect interactions between fluorescent molecules. The fluorescence lifetime of the donor fluorescent molecule is altered by FRET with the acceptor fluorescent molecule. Therefore, FLIM can analyze the degree of FRET between fluorescent molecules. FLIM-FRET has several advantages over intensity-based FRET (Bajar et al., 2016). For example, since FLIM-FRET does not require excitation of acceptor fluorescent molecules, even dim fluorescent molecules with low quantum yield can be used as acceptors (Murakoshi and Shibata, 2017). FLIM has also been used for stain-free autofluorescence imaging (Donaldson and Radotic, 2013); thus, FLIM has many potential applications.

FLIM has also been used for multi-imaging of fluorescently tagged proteins and intracellular components (Mmontova et al., 2018). However, little information is currently available about the combinations of fluorescent proteins that are suitable for the simultaneous separation based on their fluorescence lifetimes or how different fluorescence lifetimes must be in order to distinguish between different fluorescent proteins. FLIM is also expected to be useful in plant cells, which exhibit very strong autofluorescence in cell walls and chloroplasts. However, there are few examples of its application in plant cells.

Here, we demonstrate that FLIM can be used to simultaneously observe and spatially separate fluorescent proteins with very similar emission spectra and even similar fluorescence lifetimes *in vitro*. We also demonstrate that red or green fluorescent proteins fused with subcellular localization tags can be distinguished simultaneously in plant cells by FLIM.

## Results

### The signals of red fluorescent proteins are separated by FLIM *in vitro*

To evaluate whether the signals emitted by fluorescent proteins with similar emission spectra could be separated by FLIM, we selected four red fluorescent proteins: mCherry, mRFP, mApple, and tdTomato. These four proteins possess similar emission spectra that cannot be separated using conventional fluorescence filters (Table 1). We calculated the fluorescence lifetimes of each protein *in vitro* by fitting using the n-Exponential Tail Fit model. mCherry, mRFP, mApple, and tdTomato showed fluorescence lifetimes of 1.52, 1.73, 2.77, and 3.21 ns, respectively; these values are consistent with data from a previous study (Table 1). Next, we simultaneously observed two of these red fluorescent proteins *in vitro* to investigate whether signals from different fluorescent proteins with similar emission spectra could be separated and visualized based on their fluorescence lifetimes. The signal intensities of mCherry and tdTomato acquired in the range of 570–620 nm could not be distinguished (Figure 1A). However, we successfully separated their signals using FLIM, specifically using phasor plot analysis (Figure 1B, C). A phasor plot is used to visualize the distribution of fluorescence lifetimes in each pixel as a two-dimensional (2D) plot. This method allows signals from different fluorescence lifetimes to be recognized without using the complex fitting approach for multiple components (Digman et al., 2008; Malacrida et al., 2021). In addition, even though the difference between the fluorescence lifetimes of mCherry and mRFP is very small (approx. 0.2 ns, Table 1), we observed the signals from these proteins as distinct populations in the phasor plot (Figure 1D–F). Similar results were obtained using other combinations of these four proteins (Supplemental Figure S1). Furthermore, we successfully separated the signals of the four red fluorescent proteins observed at the same acquisition wavelength (570–620 nm) using fluorescence lifetime analysis (Figure 1G–I). These results indicate that fluorescence lifetime is a photophysical property of fluorescent proteins that can be used for multi-imaging analysis.

**Figure 1.**
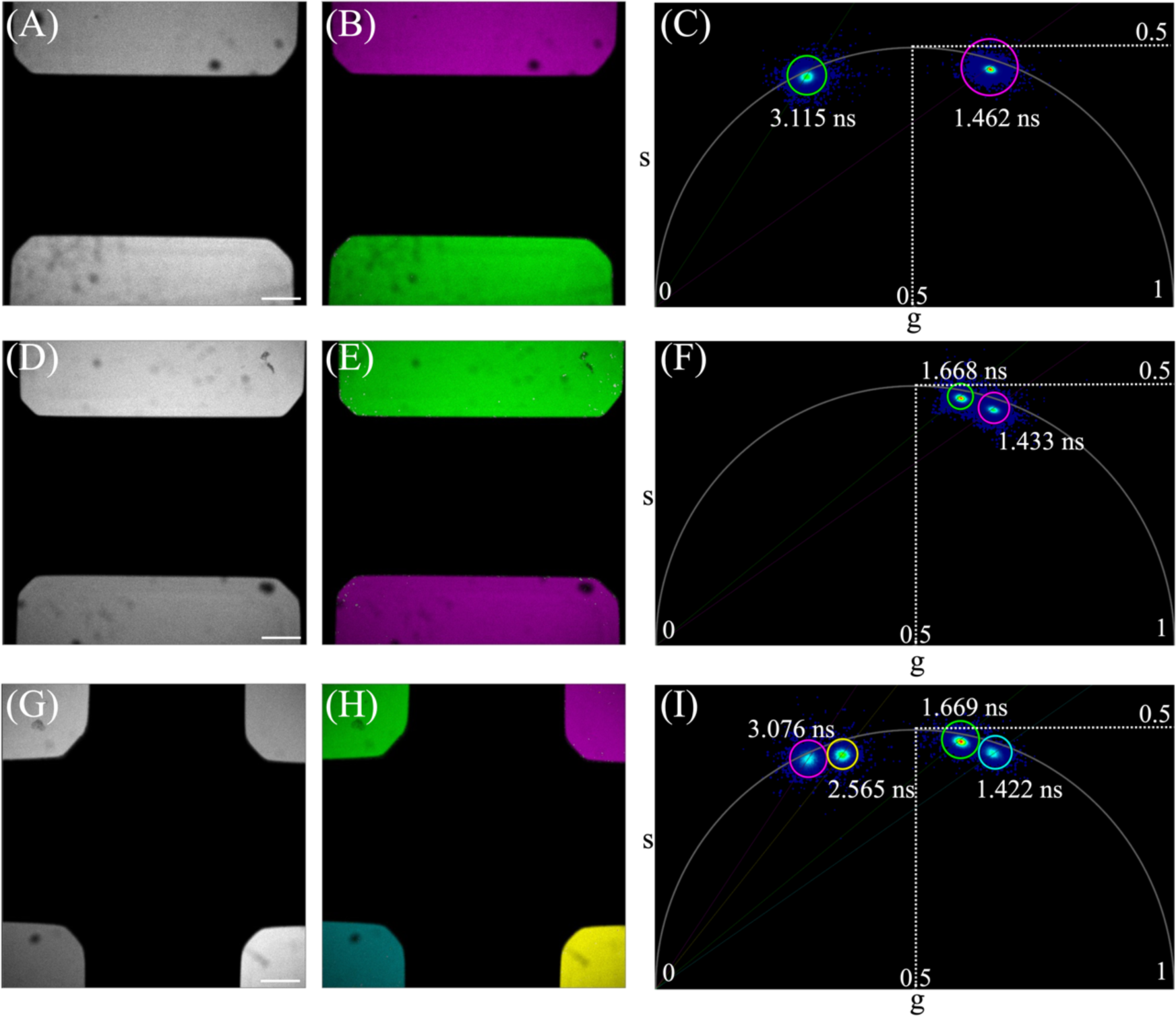
Separation of red fluorescent proteins based on their fluorescence lifetimes *in vitro*. **A–I** Fluorescence intensity images collected between 570-620 nm (A, D, G) and pseudo color images (B, E, H) based on phasor plot analysis (C, F, I) of red fluorescent proteins (mCherry, mRFP, mApple, and tdTomato). The pseudo colors in (B, E, H) correspond to the colors in the phasor plots (C, F, I). Each representative image was derived from three independent analyses. **A–C** mCherry and tdTomato are shown in magenta and green, respectively. **D–F** mCherry and mRFP are shown in magenta and green, respectively. **G–I** mCherry, mRFP, mApple, and tdTomato are shown in cyan, green, yellow, and magenta, respectively. Scale bar, 200 μm.

**Table 1.**
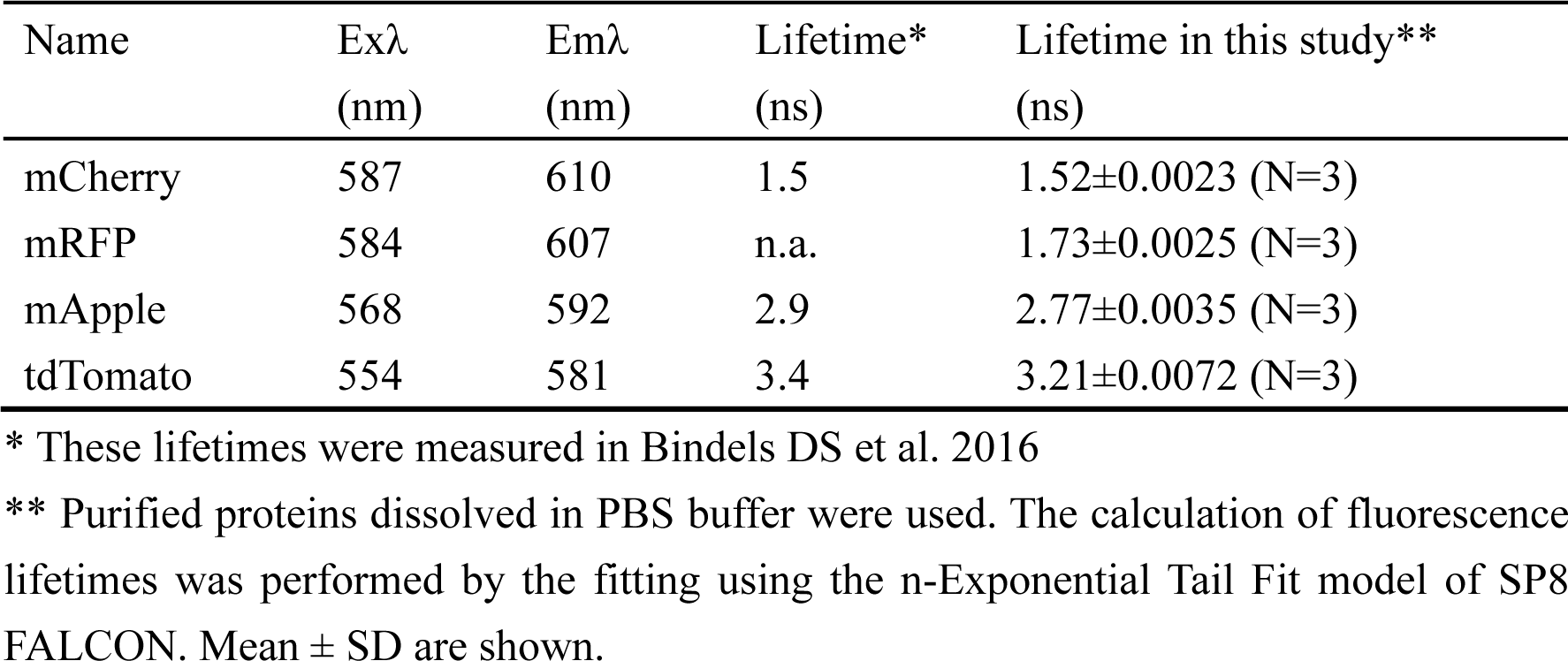
Red fluorescent proteins used for the *in vitro* experiments.

### The signals of fluorescent proteins are separated by FLIM *in vivo*

mRED7, mCherry, tagRFP-T, and mScarlet are red fluorescent proteins with very similar emission spectra but very different fluorescence lifetimes of 0.7, 1.5, 2.3, and 3.9 ns, respectively (Supplemental Table 1). To determine whether these fluorescent proteins could be simultaneously distinguished *in vivo* using FLIM, we fused each fluorescent protein with a different subcellular localization tag and transiently expressed these proteins in protonemal cells of *Physcomitrium patens* by particle bombardment. We analyzed the combination of mRED7 and mScarlet, which have the big difference in fluorescence lifetimes (approx. 3.2 ns). mRED7 fused to a nuclear-localization signal (NLS; NLS-mRED7) and mScarlet fused to a Ribulose bisphosphate carboxylase small subunit (RBCS; RBCS-mScarlet) were designed to localize in nucleus and chloroplast, respectively. The signals of these proteins were both detected in the range of 570–620 nm but could not be distinguished from each other by their intensities (Figure 2A). By contrast, we observed two distinct populations in phasor plot analysis, and these signals in the phasor plot corresponded to the intensity signals, which showed the predicted localization of each tagged fusion protein (Figure 2B, C).

**Figure 2.**
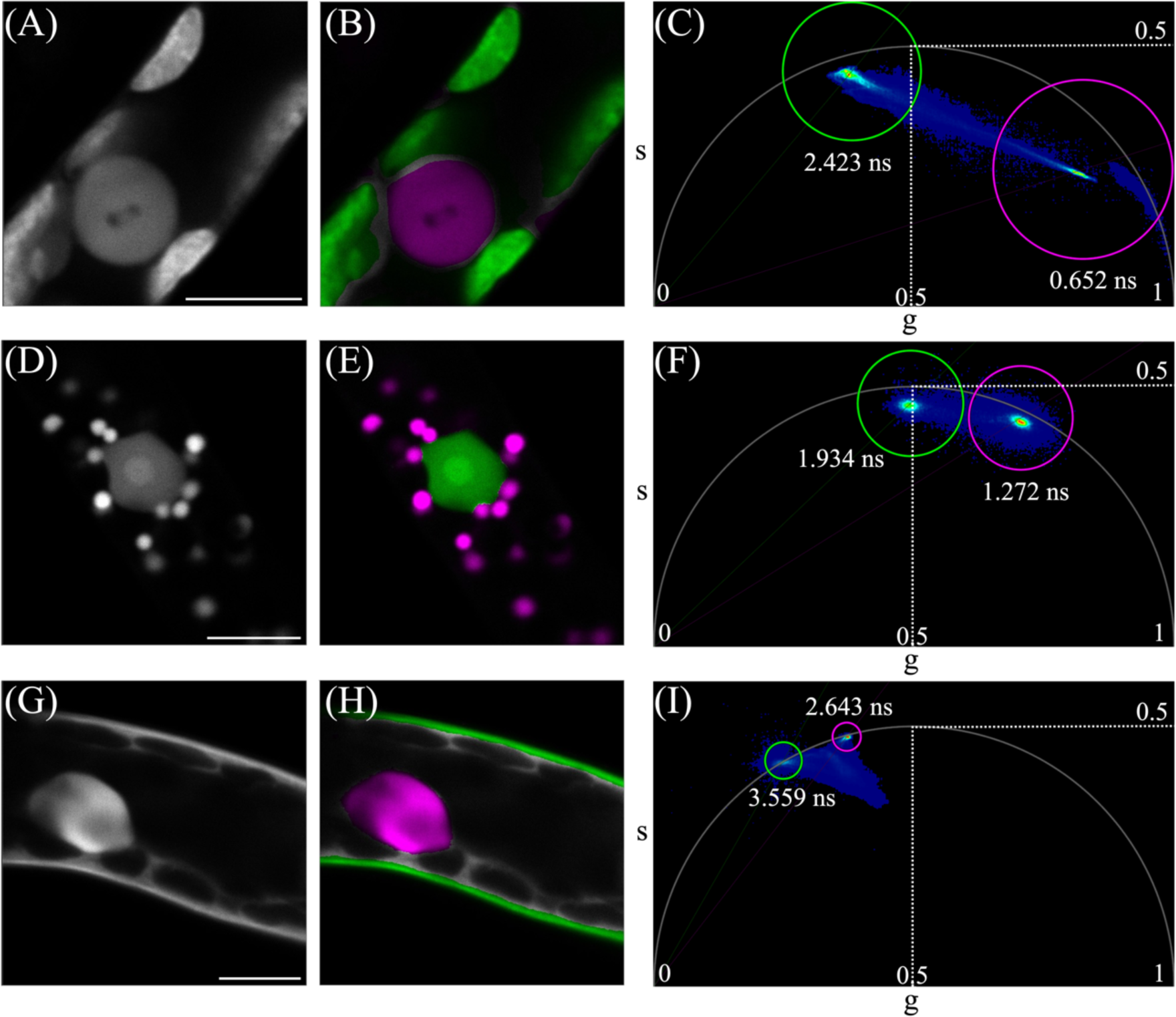
Separation of two fluorescent proteins with similar emission spectra based on their fluorescence lifetimes *in planta*. **A–I** Fluorescence intensity images collected between 570-620 nm (A, D), fluorescence intensity images collected between 490-540 nm (G) and pseudo color images (B, E, H) based on phasor plot analysis (C, F, I) of fluorescent proteins fused with a subcellular localization tag. Tag-fused fluorescent proteins were transiently expressed in protonemal cells of *P. patens* by particle bombardment. Pseudo colors in (B, E, H) correspond to the colors in the phasor plots (C, F, I). **A–C** NLS-mRED7 and RBCS-mScarlet are shown in magenta and green, respectively. The representative image set was derived from 9 independent analyses. **D–F** mCherry-SKL and NLS-tagRFP-T are shown in magenta and green, respectively. The representative image set was derived from 5 independent analyses. **G–I** NLS-GFP and NowGFP-LTI6b are shown in magenta and green, respectively. The representative image set was derived from 6 independent analyses. Scale bar, 10 μm.

We obtained similar results for the combination of the two proteins with the most similar fluorescence lifetimes: mCherry and tagRFP-T (approx. 0.8 ns difference). Although we observed mCherry-SKL (serine-lysine-leucine motif) signals in peroxisomes and NLS-tagRFP-T signals in the nucleus, these signals were both detected in the range of 570– 620 nm but could not distinguished from each other by their intensities (Figure 2D). However, we observed two different populations of signals in the phasor plot, which corresponded to the signals in the nucleus or peroxisome (Figure 2E, F). We then performed experiments using different combinations of localization tags and fluorescent proteins (NLS-mRED7 and mCherry-SKL, NLS-mRED7 and tagRFP-T-SKL, mCherry-SKL and RBCS-mScarlet, NLS-tagRFP-T and RBCS-mScarlet). For all combinations, fluorescent proteins with different fluorescence lifetimes were clearly separated using FLIM (Supplemental Figure 2A–O).

In addition to red fluorescent proteins, we also tested the green fluorescent proteins BrUSLEE, GFP, and NowGFP, with striking differences in fluorescence lifetimes (0.82, 2.8, and 5.1 ns, respectively) (Supplemental Table 1). When both NLS-GFP and NowGFP-LTI6b (LOW TEMPERATURE INDUCED GENE 6B) were transiently expressed in protonemal cells, the signal intensities of these fusion proteins acquired in the range of 490–540 nm were indistinguishable (Figure 2G). However, we observed two different populations of signals in phasor plot analysis: one population corresponded to signals in the nucleus (NLS-GFP) and the other corresponded to signals in the plasma membrane (NowGFP-LTI6b) (Figure 2H, I). Experiments using BrUSLEE-SKL and NLS-GFP or NLS-BrUSLEE and NowGFP-SKL produced similar results (Supplemental Figure 2M–R). Taken together, these results demonstrate that fluorescence lifetime is a useful tool for simultaneously observing fluorescent proteins with similar emission spectra in plant cells. Furthermore, we obtained similar results not only for different combinations of fluorescent proteins but also for combinations of different localization tags (Figure 2, Supplemental Figure 2), indicating that FLIM can separate multiple fluorescent proteins even when acquired at a single wavelength range by using different localization tags.

### Multi-imaging analysis using three fluorescent proteins *in vivo*

We next evaluated whether three fluorescent proteins can be separated *in vivo* using FLIM. When we transiently expressed mCherry-SKL, NLS-tagRFP-T, and RBCS-mScarlet in protonemal cells, the signal intensities in the peroxisome, nucleus, and chloroplast were indistinguishable, whereas we observed three distinct populations with different fluorescence lifetimes in the phasor plot (Figure 3A, C). However, we also observed signals with intermediate fluorescence lifetimes between those of mCherry and mScarlet in the boundary regions between peroxisomes and chloroplasts (Figure 3B, C). We obtained similar results in an experiment using BrUSLEE-SKL, NLS-GFP, and NowGFP-LTI6b, in which three different populations of signals in the phasor plot corresponded to the signals in the peroxisome, nucleus, and plasma membrane, respectively (Figure 3D– F). However, we also observed signals with intermediate fluorescence lifetimes in the regions between peroxisomes and the plasma membrane (Fig 3E, F). By placing a calibration bar over the signals in the phasor plot, we determined that these signals have a fluorescence lifetime intermediate between those of BrUSLEE and NowGFP (Figure 3G, H).

**Figure 3.**
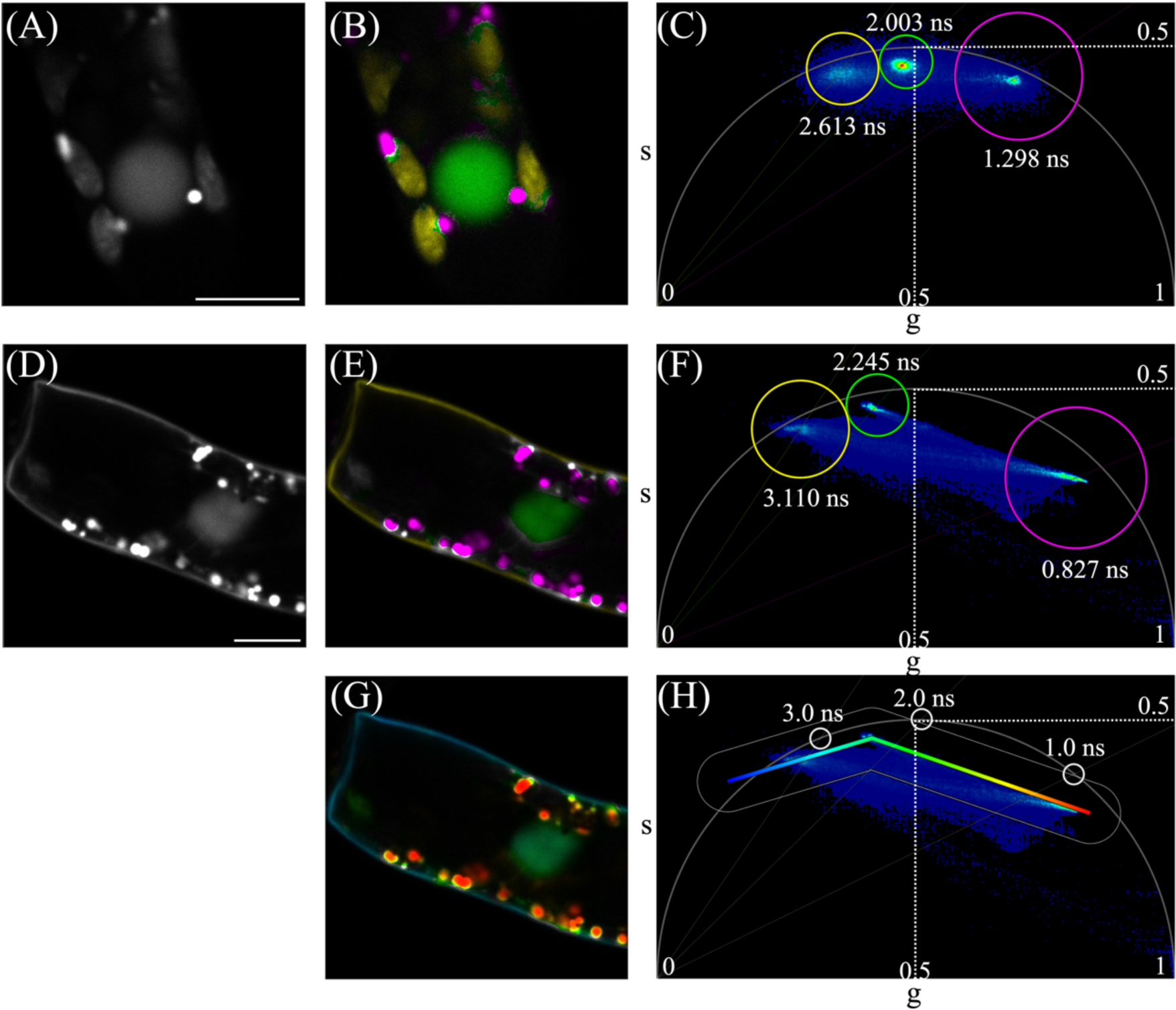
Separation of three fluorescent proteins with similar emission spectra based on their fluorescence lifetimes *in planta*. **A–H** Fluorescence intensity images collected between 570-620 nm (A), fluorescence intensity images collected between 490-540 nm (D) and pseudo color images (B, E, G) based on phasor plot analysis (C, F, H) of fluorescent proteins fused with a subcellular localization tag. Tag-fused fluorescent proteins were transiently expressed in protonemal cells of *P. patens* by particle bombardment. Pseudo colors in (B, E, G) correspond to the colors in the phasor plots (C, F, H). **A–C** mCherry-SKL, NLS-tagRFP-T, and RBCS-mScarlet are shown in magenta, green, and yellow, respectively. The representative image set was derived from 5 independent analyses. **D–H** BrUSLEE-SKL, NLS-GFP, and NowGFP-LTI6b are shown in magenta, green, and yellow, respectively (E, F). BrUSLEE-SKL, NLS-GFP, and NowGFP-LTI6b are shown in red, blue-green, and cyan, respectively (G, H). The representative image set was derived from 6 independent analyses. Scale bar, 10 μm.

## Discussion

Since confocal microscopes equipped with high-speed fluorescence lifetime devices have become commercially available, interest in using fluorescence lifetimes to achieve multi-imaging has been rapidly increasing. Particularly in the plant science field, it has been challenging to observe many molecules simultaneously in different colors due to autofluorescence from cell walls and chloroplasts, and FLIM is considered to be an effective method to solve this problem. In this study, we successfully achieved multi-imaging of fluorescent proteins with similar emission spectra using FLIM *in vitro* and *in planta* with almost no interference from autofluorescence (Figure 2, 3). In animal cultured cells, the cell cycle marker FUCCI-red was recently developed using red fluorescent proteins with different fluorescence lifetimes, mCherry and mKate2, which are fused with the cell cycle marker hCdt1 and hGem, respectively (Shirmanova et al., 2021). The FUCCI-red system can be used to analyze temporal expression variation of fluorescently labeled proteins to discriminate the cell cycle in each cell based on fluorescence lifetimes. These findings, together with our spatial discrimination of fluorescent proteins of similar emission spectra, indicate that FLIM is a powerful tool for analyzing the spatio-temporal variation of fluorescent proteins.

When molecules with different fluorescence lifetimes are present in the same location, intermediate fluorescence lifetimes would be detected depending on the ratio of their concentration. Indeed, in our fluorescence lifetime imaging, although most signals were present as distinct clusters corresponding to different fluorescence lifetimes in the phasor plot, we also observed a signal from an intermediate fluorescence lifetime in the region where the two fluorescent proteins overlapped (Figure 3). Intermediate fluorescence lifetimes were also reported in the FUCCI-red system (Shirmanova et al., 2021). In FUCCI-red, mCherry-hCdt1 and mKate2-hGem are specifically expressed during the G1 and S/G2/M phases of the cell cycle, respectively, and thus a fluorescence lifetime specific to each fluorescent protein is detected, whereas an intermediate fluorescence lifetime is observed in the G1/S phase when both fusion proteins are simultaneously expressed. These data suggest that the fluorescence lifetime signal in FLIM phasor plots provides useful information about the localization of fluorescent proteins, especially the co-localization of proteins.

Finally, we showed that FLIM with phasor plot analysis can successfully separate molecules with fluorescence lifetimes differing by as little as 0.2 ns, as long as they are spatially distinct. Due to the progress in fluorescent protein engineering, we now have numerous fluorescent proteins that can be utilized for FLIM-based multi-imaging (Goedhart et al., 2010; Mamontova et al., 2018; Mukherjee et al., 2022). In addition, several long-wavelength fluorescent proteins have been developed in recent years, which could also facilitate spectral-based multi-imaging (Chu et al., 2014; Matlashov et al., 2020). Combining spectral- and fluorescence lifetime-based imaging techniques will facilitate a more efficient, flexible analysis of the dynamics of a larger number of proteins.

## Materials and Methods

### Plant materials and culture conditions

The Cove-NIBB strain of Physcomitrium patens was used as the wild-type line (Ashton and Cove, 1977). Plants were cultured on BCDAT agar medium at 25 ℃ under continuous white light conditions. For particle bombardment, plants were blended by homogenizer and propagated on BCDAT agar medium with cellophane disc for 6-7 days.

### Construction of plasmids

Primers used in this study were listed in Supplemental Table 2. To amplify fluorescent protein sequence for construction, PIG1b-NLS-GFP-GUS, His-mCherry pCold, 35S-Lifeact-TagRFP-T pPZP211, and 35S-Lifeact-mScarlet pPZP211 plasmids were used as templates. In addition, mRED7, BrUSLEE-SKL, and NowGFP sequences were synthesized by Integrated DNA Technologies to use as PCR templates. For all construction, overexpression vector pT1O was used and linealized with *Hpa*I (New England Biolabs) for subcloning. For construction of pT1O-BrUSLEE-SKL, synthesized double strand DNA of BrUSLEE-SKL was cloned to pT1O using In-Fusion HD Cloning Kit (Takara). DNA fragments of mCherry-SKL, tagRFP-T-SKL, and NowGFP-SKL were amplified using the following primers: pT1O_mCherry_inf_F, pT1O_mCherry_SKL_inf_R, pT1O_tagRFP-T_inf_F, pT1O_tagRFP-T_SKL_R, pT1O_NowGFP_inf_F, pT1O_NowGFP_SKL_R. The amplified fragments were cloned to pT1O vector using In-Fusion for construction of pT1O-mCherry-SKL, pT1O-tagRFP-T-SKL, and pT1O-NowGFP-SKL. To generate pT1O-NLS-GFP-mRED7 and pT1O-NLS-GFP-tagRFP-T, DNA fragments of NLS-GFP, mRED7, tagRFP-T were amplified using the following primers: pT1O_NG_inf_F, NG_mRED7_inf_R, mRED7_F, pT1O_mRED7_inf_R, NLS_tagRFP_inf_R2, tagRFP-T_F, pT1O-tagRFP-T_R. The amplified fragments were cloned to pT1O vector. To generate pT1O-NLS-BrUSLEE, DNA fragments of NLS and BrUSLEE were amplified using the following primers and cloned to pT1O vector: pT1O_NG_inf_F, NLS_BruSLEE_R, BruSLEE_F, pT1O_BrUSLEE_R. For construction of RBCS-mScarlet, DNA fragments of RBCS and mScarlet were amplified using the following primers and cloned to pT1O vector: pT1O-RBCS_inf_F, RBCS_mScarlet_inf_R, mScarlet_F, pT1O-mScarlet_inf_R. DNA fragments of NowGFP and LTI6b were amplified using the following primers and cloned to pT1O vector for construction of NowGFP-LTI6b: pT1O_NowGFP_inf_F, NowGFP_LTI6b_R, LTI6b_inf_F, LTI6b_pT1O_R.

### Particle bombardment

Particle bombardment experiments were performed using a PDS-1000/He system (Bio-Rad). Gold particles with 0.6 μm diameter were washed with 100% ethanol, rinsed twice with sterilized water, and diluted with sterilized water to a concentration of 60 mg/mL. The solution (10 μL) of gold particles was mixed with 200-1500ng of plasmid DNA, and mixed with 4 μL of 0.1 M spermidine and 10 μL of 2.5 M calcium chloride. The DNA-coated gold particles were washed with 70% ethanol and 100% ethanol, and resuspended in 100% ethanol. The solution of DNA-coated gold particle was added to the macrocarrier. For bombardment, the helium gas pressure of 1100 psi and a vacuum of 25 in Hg were used. After the bombardment, moss plants were cultured for 1-2 days at 25 ℃ under continuous white light conditions to induce the expression of transgenes for FLIM analysis.

### FLIM analysis

Both intensity and fluorescence lifetime images were obtained using confocal laser scanning microscopy (TCS SP8 FALCON, Leica) equipped with a pulsed white light laser (40 or 80 MHz) and a HyD detector. HC PL APO CS2 10×/0.40 and HC PL APO CS2 20×/0.75 objective lens was used for in vitro and in vivo analysis, respectively. For in vitro analysis, four red fluorescent proteins (mCherry, mRFP, mApple, tdTomato) used in (Kurihara et al., 2021) were diluted in PBS buffer and observed using ViewPlate-1536 (PerkinElmer). To observe red fluorescent proteins, protein solution or plant cells were excited with 561 nm and their emission was acquired at 570-620 nm. To observe green fluorescent proteins, plant cells were excited with 488 nm and their emission was detected at 490-540 nm. FLIM imaging data were analyzed using phasor plot or fitting with n-Exponential Tail Fit model by LASX software.

## Acknowledgements and Funding

We thank Yoko Mizuta and Tetsuya Higashiyama for the support of particle bombardment experiment and Daisuke Kurihara for providing us with plasmid and purified fluorescent proteins. This work was supported by Yanmer Environmental Sustainability Support Association, Toyoaki Scholarship Foundation, JSPS KAKENHI Grant Number, 21K19256, 21H00364, 22H04718, JP22H04926, 22K21352, 23H04739, JST-CREST Program Grant Number JPMJCR1924, and MEXT Moonshot Program Grant Number JPMJMS2033-11 to Y.S and also supported by The Naito Science & Engineering Foundation to T.A.

---

**Supplemental Figure S1.**
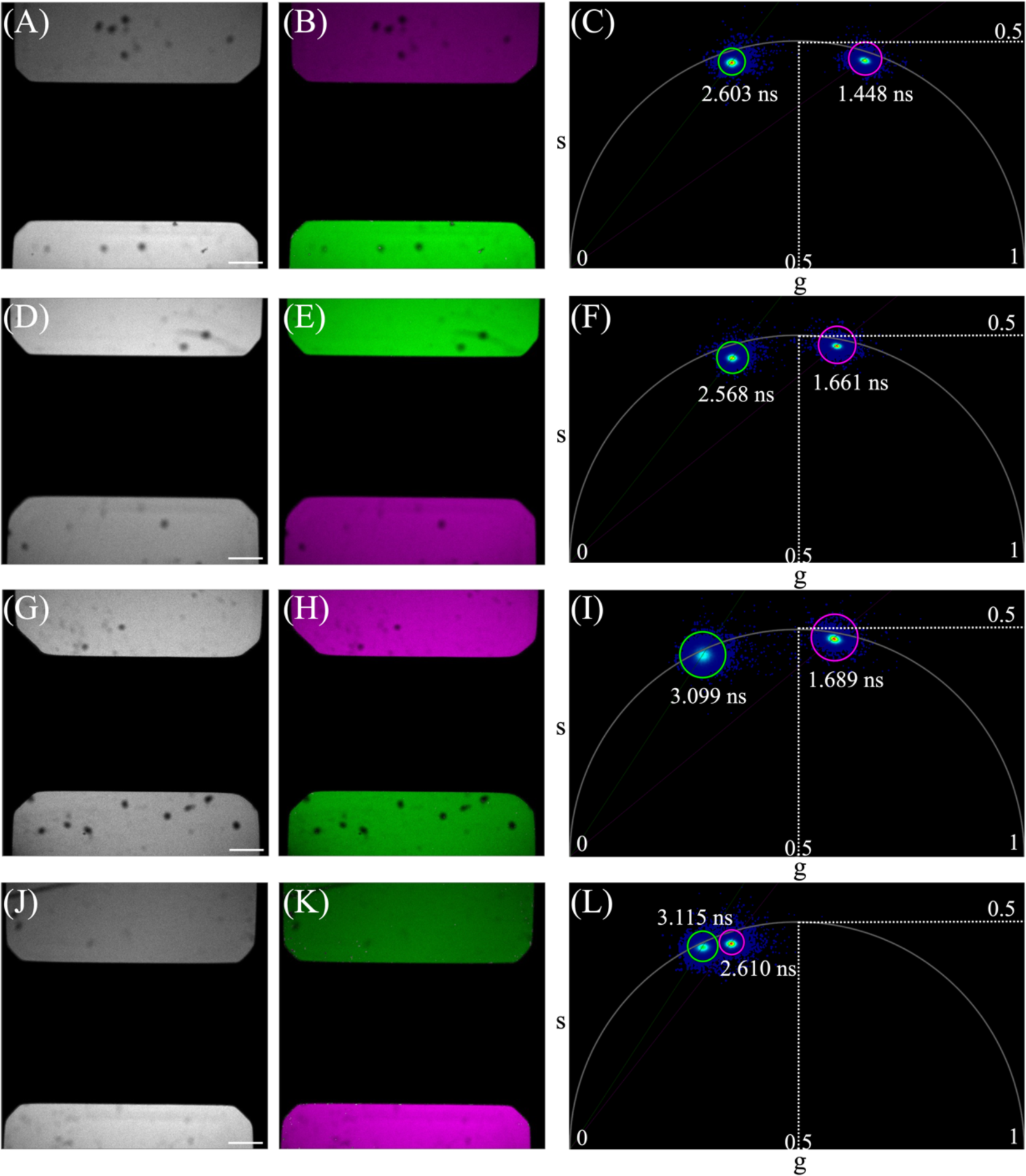
Separation of multiple red fluorescent proteins based on their fluorescence lifetimes *in vitro*. **A–L** Fluorescence intensity images collected between 570-620 nm (A, D, G, J) and pseudo color images (B, E, H, K) based on phasor plot analysis (C, F, I, L) of red fluorescent proteins (mCherry, mRFP, mApple, and tdTomato). Pseudo colors in (B, E, H, K) correspond to the colors in the phasor plot (C, F, I, L). The representative image set was derived from three independent analyses. **A–C** mCherry and mApple are shown in magenta and green, respectively. **D–F** mRFP and mApple are shown in magenta and green, respectively. **G–I** mRFP and tdTomato are shown in magenta and green, respectively. **J–L** mApple and tdTomato are shown in magenta and green, respectively. Scale bar, 200 μm.

**Supplemental Figure S2.**
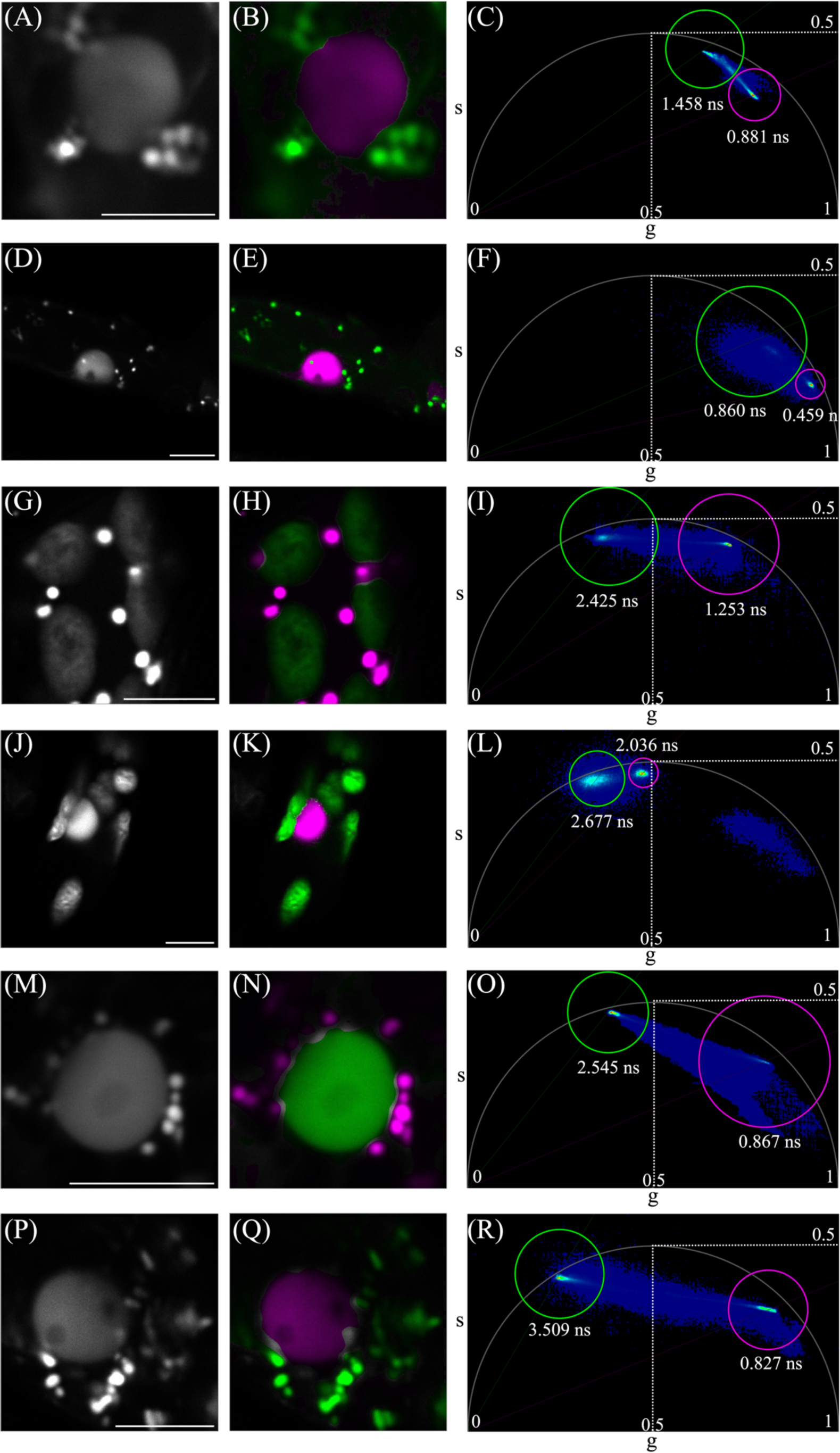
Separation of fluorescent proteins with similar emission spectra based on their fluorescence lifetimes *in planta*. **A–R** Fluorescence intensity images collected between 570-620 nm (A, D, G, J), fluorescence intensity images collected between 490-540 nm (M, P) and pseudo color images (B, E, H, K, N, Q) based on phasor plot analysis (C, F, I, L, O, R) of fluorescent proteins fused with subcellular localization tags. Tag-fused fluorescent proteins were transiently expressed in protonemal cells of *P. patens* by particle bombardment. Pseudo colors in (B, E, H, K, N, Q) correspond to the colors in the phasor plots (C, F, I, L, O, R). **A–C** NLS-mRED7 and mCherry-SKL are shown in magenta and green, respectively. The representative image set was derived from 8 independent analyses. **D–F** NLS-mRED7 and tagRFP-T-SKL are shown in magenta and green, respectively. The representative image set was derived from 2 independent analyses. **G–I** mCherry-SKL and RBCS-mScarlet are shown in magenta and green, respectively. The representative image set was derived from 6 independent analyses. **J–L** NLS-tagRFP-T and RBCS-mScarlet are shown in magenta and green, respectively. The representative image set was derived from 5 independent analyses. **M–O** BrUSLEE-SKL and NLS-GFP are shown in magenta and green, respectively. The representative image set was derived from 7 independent analyses. **P–R** NLS-BrUSLEE and NowGFP-SKL are shown in magenta and green, respectively. The representative image set was derived from 5 independent analyses. Scale bar, 10 μm.

**Supplemental Table 1.** Fluorescent proteins used for the *in vivo* experiments.

| Name | Exλ<br>(nm) | Emλ<br>(nm) | Lifetime<br>(ns) | Reference |
| --- | --- | --- | --- | --- |
| mRED7 | 589 | 606 | 0.7 | Bindels DS et al. 2016 |
| mCherry | 587 | 610 | 1.5 | Bindels DS et al. 2016 |
| tagRFP-T | 555 | 584 | 2.3 | Bindels DS et al. 2016 |
| mScarlet | 569 | 594 | 3.9 | Bindels DS et al. 2016 |
| BrUSLEE | 487 | 509 | 0.82 | Mamontova et al. 2018 |
| GFP | 488 | 507 | 2.8 | Mamontova et al. 2018 |
| NowGFP | 494 | 502 | 5.1 | Sarkisyan et al. 2015 |

**Supplemental Table 2.**
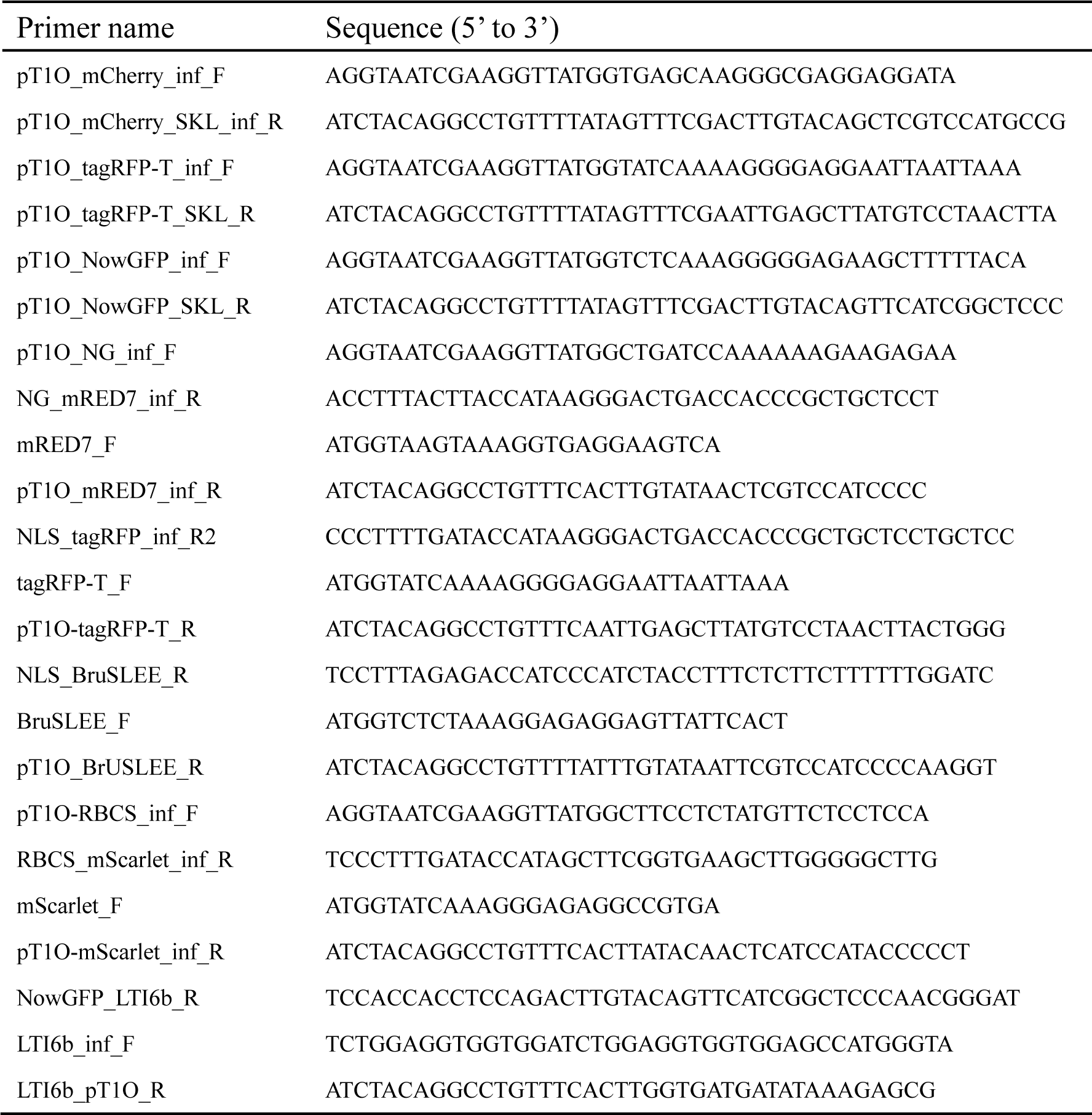
Primers used in this study.

